# Characterizing and interpreting the influence of internal variables on sensory activity

**DOI:** 10.1101/114439

**Authors:** Richard D. Lange, Ralf M. Haefner

## Abstract

The concept of a tuning curve has been central for our understanding of how the responses of cortical neurons depend on external stimuli. Here, we describe how the influence of unobserved internal variables on sensory responses, in particular correlated neural variability, can be understood in a similar framework. We suggest that this will lead to deeper insights into the relationship between stimulus, sensory responses, and behavior. We review related recent work and discuss its implication for distinguishing feedforward from feedback influences on sensory responses, and for the information contained in those responses.

**Highlights:**

- Re-interpretation of neural correlations in terms of internal variables…
- …can clarify whether they limit or enhance information
- Influence of internal variables can be captured by interpretable ‘tuning functions’
- Estimation of both internal variables and tuning possible from population recordings

## Introduction

A large part of cortical function can be characterized as transforming sensory inputs to behavioral outputs. While this transformation is conceptually unidirectional, the anatomical structures implementing it are largely bidirectional [1]. In this review, we concentrate on discussing progress in understanding ‘top-down,’ or feedback signals (FB) that influence the responses of sensory neurons. While the correlated variability of neural responses has traditionally been interpreted primarily as noise and in terms of local network connectivity or limited input information (reviewed in [2]), it may also arise from variability in unobserved internal variables projecting back to sensory areas [3, 4^••^, 5^••^]. We summarize and compare experimental techniques for inferring those internal variables from sensory population recordings, and show that fluctuating internal variables offer simple and intuitive accounts of many empirical results. While our focus here is on empirical studies of early visual areas, we expect the discussed techniques and computational framework to also be useful in understanding neural responses in other sensory or motor areas [6•].

The brain’s representation of sensory inputs and transformation into behavioral outputs can be understood on at least two levels. On the first one, we would like to understand how a stimulus **(S)** influences neural responses **(r)**, and how these responses influence behavior **(B)**. On a more abstract level, one can try to understand how a stimulus affects abstract internal variables **(I)**, and how these variables are linked to behavior (Figure 1). While these two levels are clearly related, the representation of abstract internal variables is generally unclear, and it is in principle only possible to directly observe the corresponding neural responses; internal variables are only accessible by their correlation with one or more of the observable quantities, **S, B**, and (a subset of) r (Figure1).

**Figure 1:**
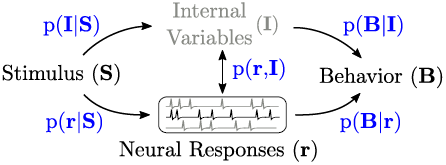
A computational-level description of the brain (upper path) may invoke abstract internal variables **(I)** that govern behavior **(B)** and are influenced by stimuli **(S)**. On a neurophysiological level, we seek a similar description in terms of the responses **(r)** of populations of neurons (lower path) [7]. It is often useful to mix levels of abstraction, for example modeling the relationship between attention (an abstract state, or internal variable) and neural responses. Studying these relationships is limited by what quantities may be directly observed; **I** is not directly observable, and r is only partially observable (indicated in gray). Experiments in which an estimate of **I** is available from an observable correlate (**S** or **B**) lend themselves to *conditional* approaches, estimating p(**r**|**I**). *Unsupervised* approaches model the relationship between r and I, estimating both the instantaneous value of an internal variable and its influence on responses given only observations of neural responses p**(r)**.

Mathematically, relationships between these variables may be framed either as conditional or as joint probability distributions. Conditional probability distributions are commonly used when one of the variables is known, or if a causal role of this variable is hypothesized, for example the stimulus-conditioned response distribution, the mean of which defines the tuning function with respect to that stimulus. Thinking in terms of joint probability distributions, on the other hand, is useful to characterize a wider range of relationships that are not necessarily causal, and where neither variables needs to be under the control of the experimenter, or even be observed. In the following, we review existing work in the context of these two approaches and relate them to attempts to distinguish feedforward from feedback signals in the brain. In particular, we describe how the concept of a ‘tuning curve’ can be extended to capture the relationship between neural responses and internal variables (Box 1), and further show that this yields simple and intuitive descriptions of correlated noise in sensory areas (Box 2).

## 1 Inferring internal variables and their influence on neural responses

Since a subject’s internal variables are not observable directly, the traditional approach to measuring their impact on neural responses has been to exploit their relationship to observed quantities like stimulus and behavior. For example, training an animal in an attention task induces a relationship between an internal variable (‘attentional state’) and an external sensory signal (‘attention cue’)[3], where improved performance on a task is taken as an indicator for successful manipulation of attentional state [8^••^].

The first limitation of this approach is the fact that there usually remains significant uncertainty about the relationship between observables and internal variables. For instance, even though an attention cue may only have two states, the animal’s true attentional state is likely non-binary and more variable. This limitation can be overcome by bootstrapping from a binary cue to a continuous internal representation and by making additional assumptions about the relationship between the internal variable and the measured neural responses. Assuming linearity, for instance, allows one to define an ‘axis’ connecting the average neural responses conditioned on the two possible cues, along which neural responses will vary as the internal variable varies (Box 1). Projecting the neural activity on a single trial onto this axis, allows one to estimate an intermediate value of the internal variable [8^••^]. Cohen and Maunsell [8^••^, 9, 10, 11] indeed showed that the estimates obtained using this approach in a change-detection task while recording from neurons in area V4, predicted behavioral performance (accuracy) on single trials. An analogous approach can be taken to infer a ‘choice axis’ [12, 13] which is closely related to measuring choice probabilities across a population of sensory neurons [14, 15]. The projection of the population activity on a single trial onto this choice axis yields an estimate of a decision-related variable. If correct, this projection will correlate with behavioral measures of confidence or reaction time. Such an approach to inferring the state of an internal variable might be called *conditional* since it relies on an observable correlate of the variable of interest (Fig. 1).

The second limitation, which the conditional approach cannot overcome, is the fact that internal variables are surely richer than the behavioral output in perceptual decision-making tasks (especially when behavior is limited, as in discrimination tasks), or the number of trainable cues. For instance, ‘attention’ is likely to be a higher-dimensional variable incorporating different location(s), feature(s), etc. to which the animal is attending [9, 16•]. Equally, the brain is likely to form variable internal beliefs beyond those related to the behavioral report in a particular task. One way to overcome this limitation is to exploit the statistical structure induced in population responses by variability in those internal variables. Fluctuations in an (unobserved) internal variable increases (observable) response variability in a direction that can be thought of as the neural responses’ ‘tuning’ to that variable (Box 1, Figure 2). Different internal variables all leave their own signatures that superpose (Figure 2b, Boxes 1 + 2). By fitting the *joint* probability distribution p(r, I), it is possible to infer both the internal (‘latent’) variables as well as their influence on the responses from observations of the responses alone alone, in an unsupervised way [6•, 17, 18]. A well-known example of this approach is Factor Analysis, with each ‘factor’ corresponding to an internal variable in this context. The ‘factor loadings’, i.e. each factor’s influence of neural responses, corresponds to an ‘axis’ recovered in the conditional approach above, and can in turn be used to infer estimates of the corresponding internal variable over time [4^••^], or on a trial-by-trial basis [5^••^]. ‘Demixed PCA’ [19] presents a promising mixed approach, incorporating hypotheses about multiple internal variables into jointly fitting p(r, I). Knowledge of the inferred variables’ values and their corresponding ‘tuning curves’ may allow a functional interpretation of them, e.g. as the influence of anesthesia [4^••^], attentional state [8^••^, 20, 21, 22], motor activity [23], or in terms of their sensory representation, e.g. as beliefs about the outside world [15, 24, 25^••^, 26, 27, 28]. Even though this approach requires large neural data sets (but not prohibitively large [29]), we believe that it will be an extremely productive one (see Box 2 for an illustration).

**Figure 2:**
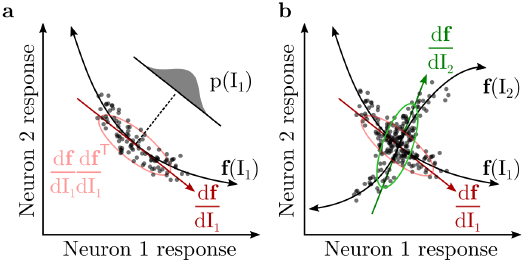
a: The population tuning of two neurons to some variable, f(I_1_), entails responses spread along f when I_1_ varies according to p(I_1_) (each dot indicates the population response on a single trial) [31•]. The linear approximation to f models this as co-variability along 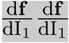(ellipse), which we call the ‘signature’ of I_1_ in this population. **b:** Variability in a second, independent internal variable, I_2_, sums (equation (2)). Given certain assumptions and enough data, unsupervised methods are able to infer I_1_, I_2_, and f(I_1_, I_2_) from observations of responses alone.

#### Box 1: How fluctuations in internal variables induces correlated variability in neural responses

A neuron’s response can be modeled as a function of both *internal* and *external* variables:

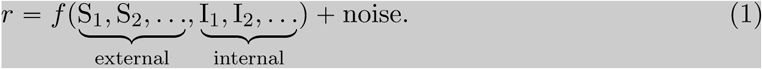

Locally, this *tuning function* can be approximated linearly by its slope with respect to each internal and external variable. Any variability in its arguments will induce variability in the neural response [3, 30]. Across a neural population, **r** = (*r*_1_, *r*_2_, ..), this variability will be correlated with the following response covariance^†^

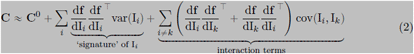

where **C**^0^ is ‘intrinsic’ covariance (i.e. the covariance of the ‘noise’ term that is not attributed to **S** or **I**). The ‘interaction terms’ appear whenever two states are not independent, though independence might be assumed for simplicity. Terms due to variability in S have been omitted to focus on so-called ‘noise correlations,’ or correlated variability while keeping the stimulus constant.

By analyzing the trial-by-trial variability in the responses of a population of neurons (illustrated as dots in Figure 2), we may recover df/dI, and from this we learn about the internal variables modulating neural responses.

#### Box 2: Case Study: Cohen and Newsome (2008) [32^••^,]

Following Box 1 we show how the empirical correlations reported in [32^••^,] can be understood in terms of the (linear approximation of) tuning of the neural responses to two internal variables. The authors measured pairwise noise correlations between neurons in area MT during a motion discrimination task, where the discriminated directions changed from trial to trial, but were always separated by 180°. Comparing correlations for the same zero-signal external stimulus, the authors found that noise correlations changed depending on whether the two neurons either supported the same decision, or opposite decisions (‘same pool’ and ‘different pool’ respectively, Figure 3c).

**Figure 3:**
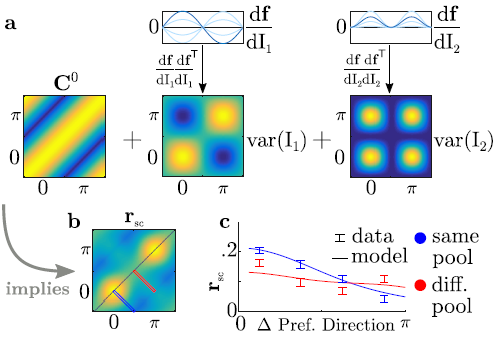
a: Visualization of equation (2) (Box 1), plots in a-b shown with respect to neurons’ preferred motion direction relative to the task. Assuming limited range structure for C^0^ and two independently-fluctuating internal variables that act selectively on the task-relevant directions 0 and *π*. b: The sum of covariance matrices in a implies the correlation (r_sc_) structure shown here. Red and blue rectangles indicate slices shown in the next panel. c: Cohen and Newsome (2008) [32^••^,] measured pairwise correlations as a function of the difference in the pair’s preferred directions in two (‘same pool’, ‘different pool’) conditions. Data replotted as error bars and compared to model (lines). Note that in the absence of I_1_ and I_2_, the two conditions would have identical correlation structure.

The data are well-explained by two independently varying internal variables, I_1_ and I_2_ that influence neural responses according to the tuning functions, df/dI_1_ and df/dI_2_, shown in Figure 3a. I_1_ increases the gain to neurons that prefer one of the task-relevant motion directions while suppressing responses of neurons that prefer the other direction, or vice versa. I_2_ varies independently of I_1_, increasing the gain of all neurons selective to either task-relevant motion direction together. While these factors were hypothesized post-hoc by the authors, they can in principle be inferred directly from population recordings using the *unsupervised* methods discussed in the text. Characterizing neural variability in these terms promises deeper insights into whether any of the internal variables are best understood as graded beliefs [26, 27, 28] (if they are correlated with choice and confidence), categorical actions [33] (if they are correlated with choice but not confidence), attentional states [8^••^, 20, 21] (if they are correlated with performance), or general network states [4^••^]. Here, the authors characterized the two variables as fluctuating ‘attention’ on one versus the other motion direction (I_1_), and fluctuating ‘attention’ to the task in general (I_2_) [32^••^].

## 2 Distinguishing feedforward from feedback influences

Two non-causal lines of evidence can help distinguish between FF and FB influences: (1) comparing measured responses to the predictions from a sufficiently powerful FF model and ascribing residuals to FB influences, and (2) comparing neural responses to identical stimuli for different induced states of internal variables under the *assumption* that they are represented by neurons downstream of those that are being observed (Figure 4). Note that we refer here only to the functional definition of FB as ‘top-down’ signals, and that FB signals from downstream neurons may themselves involve not just pure cortico-cortico FB pathways but also anatomically FF pathways [34], for instance via a thalamocortical loop [35, 36].

**Figure 4:**
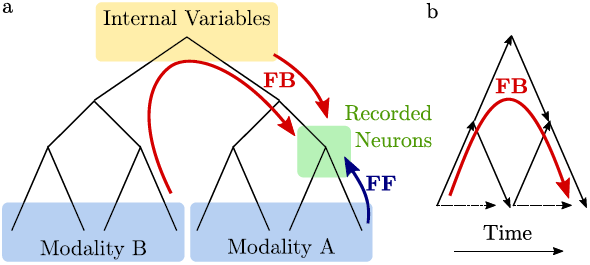
Cartoon of the functional relationships involved in common FB, or ‘top-down’ signals. a: FB through hierarchical structure: from the perspective of a recorded area (green), e.g. area V4 in the visual modality, lower areas within that modality are considered FF, and influences from other modalities like audition are considered FB. FB signals may also arise from intrinsically generated abstract internal variables (yellow), such as motivation, attention, or beliefs. b: FB through time: dependencies on external inputs at different times, sufficiently long in the past such that ‘memory’ about them within the FF pathway can be excluded, may also be considered FB.

Nienborg and Cumming (2009) [37•] took the first of these approaches. Showing that the correlation of neural responses and behavior, p**(r, B)**, had a different time course from the correlation between stimulus and behavior, p**(S, B)**, they concluded that p**(r, B)** must at least in part be due to FB signals.

Most studies taking the second approach manipulate the attentional state of the subject, e.g. by presenting a pre-trial cue (Figure 4b), and have been extensively reviewed elsewhere [38]. A newer approach relying on population recordings is to analyze the co-variability of the neural responses and determine whether they contain task-dependent components. Under the critical assumption that task-learning, or task-switching, does not alter the FF connectivity or recurrent connectivity, one can then conclude that any such task-dependent component must be due to FB signals. The underlying assumption is more likely to be true for early sensory areas than higher levels of sensory processing where learning-induced plasticity has observed to be stronger [39]. The assumption is also more likely to be true for task-switches across shorter time scales, e.g. from trial to trial (seconds) rather than days or weeks over which relevant changes of the FF pathway become more likely. Both Cohen and Newsome (2008) [32^••^] (see Box 2) and Bondy and Cumming (2016) [25^••^] have taken this approach comparing changes across trials in area MT responses, and changes separated by several days in area V1 responses, respectively. Even though Rabinowitz et al (2015) [5^••^] did not explicitly vary the task, a similar logic can be applied: because the ‘tuning curve’ associated with one of the main latent variables driving trial-to-trial variability has a task-specific shape, it has to be due to an internal variable affecting neural responses through FB signals.

In all of these studies, training on a task induces a task-specific relationship between internal variables and sensory responses; this relationship is ascribed to FB under the assumption that those variables are represented by a ‘downstream’ area.

## 3 Implication for the information in sensory responses

As shown in Box 1, a fluctuating internal variable induces correlated variability in a population proportional to the population’s ‘tuning’ to that variable. When ‘tuning’ to this internal variable, df/dI, is in the same direction as the population’s tuning to the stimulus, df/dS, then variability in **I** has the *potential* to limit information about S, since it would result in changes in the population response that are indistinguishable from changes to the stimulus itself [31•]. However, if the internal variable covaries with the stimulus in a useful way, then it may enhance the information the population carries about S. For example, random trial-to-trial fluctuations in attention can only reduce information [16•] (because they are uncorrelated with the stimulus), while ‘justified’ beliefs about the stimulus that vary trial-to-trial do not [15, 27].

Hence the question of whether top-down sources of correlated variability are helpful or harmful for population coding depends not just on the ‘shape’ of the induced correlations, but more generally on whether the internal variables underlying them carry information about the stimulus.

## 4 Conclusion

The influence of internal variables on sensory responses can be characterized by tuning functions in analogy to the influence of the external stimulus on neural responses. Uncontrolled fluctuations in those variables leaves characteristic signatures in the statistics of sensory responses that can in principle be exploited to infer the state of those variables and variable-specific tuning functions. Interpreting tuning to internal variables and inferring their states allows us to ask new questions (Box 2): Which internal variables contain information about the stimulus, and which about the choice [13]? Which variables are correlated with overall performance in the task? Do they correspond to variables in a computational model of the brain performing the task? Answers to these questions promise deeper insights into the relationship between correlated variability and stimulus information in neural populations, and may allow us to bridge Marr’s level of hierarchy between computational models and their neural implementation [7].

## Acknowlegdements

We thank Alexander Ecker, Ruben Moreno-Bote, Daniel Chicharro, Jacob Yates, Greg DeAngelis, and Aki Anzai for feedback on the manuscript.

## Footnotes

† Correlations between **I** and r considered here may reflect causation from **I** to r, or common input to both.

